# Allostery links hACE2 binding, pan-variant neutralisation and helical extension in the SARS-CoV-2 Spike protein

**DOI:** 10.1101/2025.01.28.635280

**Authors:** Alice Colyer, Esther Wolf, Cristina Lento, Mart Ustav, Adnan Sljoka, Derek J. Wilson

**Author notes:** **Correspondence to Derek J. Wilson**. **Contributed equally** (Alice Colyer, Esther Wolf).

## Abstract

The SARS-CoV-2 spike protein is highly antigenic, with epitopes in three distinct regions of the receptor binding domain (RBD) alone that have known mechanisms of neutralization. In previous work, we predicted a fourth RBD epitope based on allosteric conformational perturbations measured by hydrogen-deuterium exchange mass spectrometry (HDX-MS) upon complexation with the canonical spike protein target, human angiotensin-converting enzyme 2 (hACE2). We subsequently identified a pan-neutralizing antibody (ICO-hu104) with the predicted epitope, however, as the epitope was somewhat distant from the hACE2 binding interface, and our previous work limited to the spike RBD, the neutralization mechanism was unclear. Using HDX-MS, we investigated the binding of ICO-hu104 to the full-length SARS-CoV-2 spike protein from Wuhan, Delta and Omicron variants. We demonstrate that binding of ICO-hu104 at its epitope results in an increase in deuterium uptake in the distant HR1 domain for variants of concern, which in a biological context could be indicative of destabilisation of the helices within this region, promoting S1 shedding or failure of helical extension during S2-mediated fusion. This is supported by our computational modelling, highlighting propagation of allosteric effects to the S2 coiled-coil region. Collectively, this work demonstrates an alternative neutralization mechanism for ICO-hu104 which is distinct from its first-generation predecessors and thus opens alternative avenues targeting non-RBD epitopes through assessment of allosteric perturbations.

**Highlights:**

- HDX-MS reveals decreased deuterium uptake within the HR1 region of S2 for SARS-CoV-2 spike protein for variants of concern when bound to ICO-hu104.
- Computational modelling validates high allosteric coupling between ICO-hu104 epitope and HR1 region.
- Suggests an alternative neutralization mechanism from its predecessor ICO-hu23, whereby destabilisation of helices within the HR1 region promotes S1 shedding and/or failure of helical extension during S2-mediated fusion.

**Graphical abstract:** 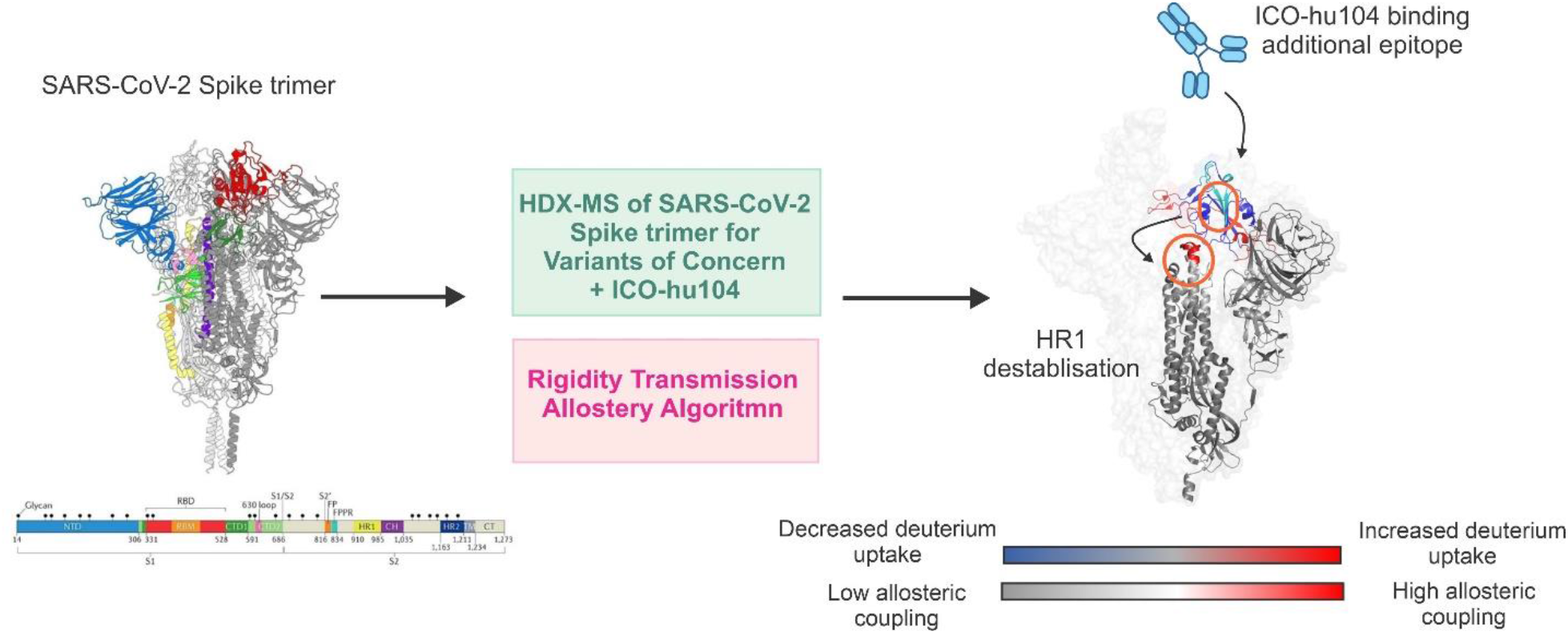

## Introduction

The development of antibody-based therapeutics for SARS-CoV-2 remains an ongoing endeavour particularly to combat viral variants of concern (VoC) which still pose a threat to public health systems [1,2]. The SARS-CoV-2 virion is comprised of four structural proteins (nucleocapsid, membrane, envelope and spike) in addition to 16 non-structural proteins (NSPs) and 6 accessory proteins **(Figure 1A)** [3]. The spike glycoprotein is a homotrimeric assembly that extends from the surface of the virion. Each monomeric subunit is subdivided into two regions, termed S1 and S2, delineated by furin protease cleavage in the Golgi apparatus during viral maturation in infected cells [4]. S1 comprises the N-terminal domain (NTD) and the RBD, which mediates attachment of the spike protein to the hACE2 receptor in the surface of the host cell [5]. S2 is composed of an S2’ cleavage site, which exposes a fusion peptide facilitating membrane fusion, in addition to a transmembrane domain and two heptad repeats (HR1 and HR2), which form a six-helical coiled coil **(Figure 1B)** [6]. Infection begins by binding of the spike protein at the RBD *via* hACE2, mediating endocytosis (specifically *via* the receptor binding motif [RBM] at residues 437-508 interacting with the hACE2 N-terminal helix) [4]. The viral particle is engulfed into an endosome, where the non-covalently attached S1 detaches and S2 extends to expose the S2’ cleavage site, being cleaved by host cell proteases and initiating host-virion membrane fusion, allowing the positive sense, single stranded viral mRNA genome to enter the host cell [4].

**Figure 1.**
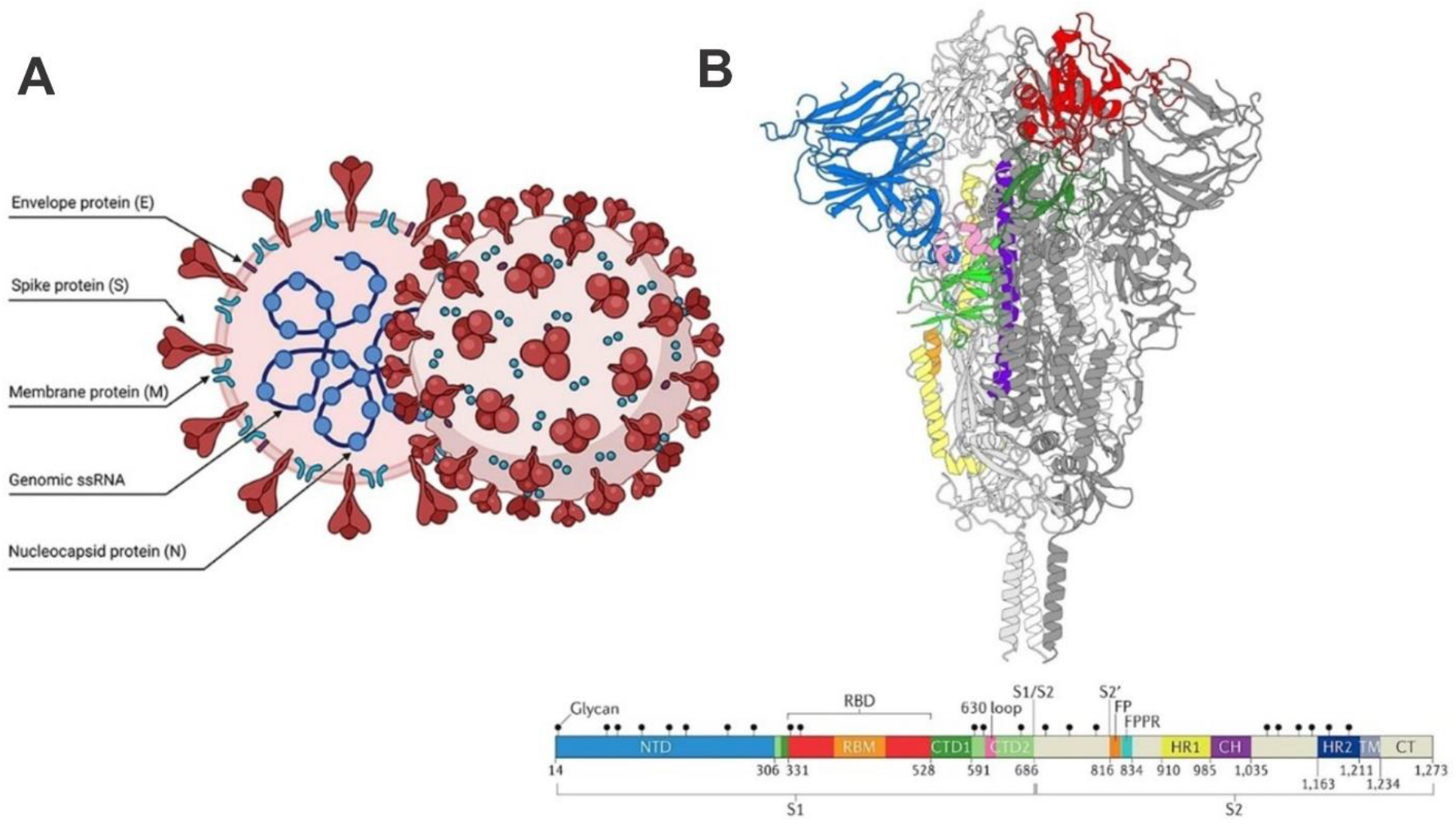
SARS-CoV-2 Virion and Spike Protein. (A) The enveloped viral particle has several structural proteins which enclose genomic positive sense, single-stranded RNA: envelope (E), spike (S), membrane (M), and nucleocapsid (N). (B) The spike protein has distinct domains, including a receptor binding domain (RBD, green) and an N-terminal domain (NTD, blue). It is cleaved into S1 and S2 during virion formation at the S1/S2 site. S1 and S2 are noncovalently bound upon hACE2 recognition. (a) and (b) were reproduced with permission from Pizzato et al. (2022) (Open access under CC BY 4.0 DEED) and Jackson et al. (2021) [4,7].

Initial antibody-based therapeutics were designed to disrupt the interaction between the spike glycoprotein and hACE2 receptor *via* the S1 subunit / RBD [8–10]. In addition, infection of SARS-CoV-2 is known to promote conformational switching within the RBD of the spike protein, alternating between an ‘up’ (receptor accessible) or ‘down’ receptor inaccessible conformation [11]. Understanding how therapeutics target either the ‘up’ or ‘down’ conformations of the SARS-CoV-2 spike protein is essential in order to progress and compliment therapeutic design, especially due to the rapid rate at which the spike protein accumulates mutations within the RBD, which can quickly render therapeutics ineffective.

Our previous HDX-MS work predicted three potentially neutralising epitopes located within the RBD [12]. Whilst the neutralisation mechanism of two of these epitopes had previously been documented, revealing direct interference of hACE2 binding [13], the third epitope did not appear to have any known antibodies and therefore no neutralization mechanism identified.

Subsequently, we identified a pan-neutralising antibody developed by ICOSAGEN (ICO-hu104) that binds to this third epitope, which is at a location somewhat removed from the hACE2 binding site within an exposed β-sheet region (residues 433-443) [12,14]. To explore this interaction, we have utilised ΔHDX-MS; an approach that has proven exceptionally powerful for epitope mapping and for the determination of neutralization mechanism across many systems [15–18]. Furthermore, high affinity previously reported between ICO-hu104 and its RBD epitope enables orthosteric and/ or allosteric perturbations to be readily observed upon binding of ICO-hu104 to full-length spike protein by ΔHDX-MS. Together, our results highlight how HDX-MS can be applied to uncover a possible neutralization mechanism of the SARS-CoV-2 spike protein for VoC revealing allosteric unravelling of HR1 within the S2 region, preventing membrane fusion.

## Materials and methods

### Protein expression and purification

Proteins were expressed in CHO cell lines and purified using Ni-NTA affinity chromatography by our coauthors, Hermet et al. (2023),1 including the Wuhan RBD (res. 319-541, Icosagen CAT#: P-307-100), Omicron RBD (B.1.1.529, res. 319-537, Icosagen CAT#: P-3607-100), Trimeric Spike Omicron (B.1.1.529, res. 14-1211, Icosagen CAT#: P-369-100), Trimeric Spike Delta (B.1.617.2, res. 14-1211, Icosagen CAT#: P-353-100), and Trimeric Spike Wuhan (res. 14-1211, Icosagen CAT#: P-309-100). The trimers had inactivated S1/S2 cleavage site (GSAS) and a C-terminal T4 fibritin trimerization sequence to improve in vitro stabilization of the intact trimer.

### HDX-MS epitope mapping

Hydrogen-deuterium exchange was carried out using the commercial Waters HDX system. Briefly: samples were mixed and injected in a labeling time-randomized order by a PAL3 Autosampler followed by UPLC separation and mass spectrometry analysis using a Waters M-Class ACQUITY UPLC with peptides detected on a Waters Select Series Cyclic IMS Mass Spectrometer. Labeling times of 1, 10, and 60 min at RT were used, followed by quenching (7.5 M guanidine hydrochloride and 0.5M TCEP) at 0°C and digestion using an Enzymate BEH Pepsin column. The peptides were reverse-phase separated using an ACQUITY UPLC BEH C18 column. Peptide identification was carried out using HDMSe with CID fragmentation in the transfer cell and ProteinLynx Global Server (PLGS), followed by HDX analysis using DynamX. Cumulative Woods plots (summed across all HDX-MS time points) were generated using Deuteros [19], identifying statistically significant peptides with an increase or decrease in deuterium uptake with an applied confidence interval of 98%. Structures were visualized using PyMOL 2.5.

### Computational allostery modelling

Starting with the trimeric Wuhan and Omicron Spike protein structures (PDB 7DDN and PDB 7DGW), we applied Rigidity Transmission Allostery (RTA) algorithms [20,20,21] to predict potential allosteric effects on spike protein due to perturbations of the epitope residues identified by HDX-MS. RTA method and algorithms are based on mathematical rigidity theory which have been extensively applied in allostery studies of GPCRs, enzymes, DNA-binding proteins and many other systems [22–25]. RTA algorithms, whose details have been previously described, provides a mechanistic model and prediction of allosteric coupling in protein structures. In short, RTA predicts if local perturbations of rigidity and conformational degrees of freedom propagate and modify rigidity and conformational degrees of freedom of distant sites across proteins and their complexes.

We used the FIRST software [26] (Floppy Inclusions and Rigid Substructure Topography) to construct a constrained network (graph), where nodes represent atoms and edges represent covalent and non-covalent interactions (hydrogen bonds, hydrophobic interactions etc.). Hydrogen bonds are ranked based on their individual energy strengths. We perturbed the rigidity of the epitope residues identified by HDX (470-512 for ICO-hu23-Wuhan and 433-441 and 496-512 for ICO-hu104-Wuhan and 433-443 and 497-510 ICO-hu104-Omicron) and then applied the RTA algorithms to extract the conformational degrees of freedom for other residues, before and after the perturbation of the epitope. To capture allosteric effects, we are interested in the change (i.e. transmission) in the available conformational degree of freedom after perturbation. The “degree of freedom transmission” (i.e., allosteric transmission) was quantified for individual residues and to visualize the allosteric networks (pathways), residues are color-coded on the 3D structure based on the strength of their allosteric transmission.

## Results

### Effects of ICO-hu23 and ICO-104 on full-length spike protein

Briefly, ΔHDX-MS was performed on ICO-hu23 upon complexation with the full-length spike proteins from Wuhan and ICO-hu104 with Wuhan, Omicron and Delta. ΔHDX-MS of the unbound state was performed at 15 μM and the bound state at a 1:1 equimolar concentration, with labelling performed at 5, 10 and 30 minutes. The exchange reaction was quenched by lowering of the pH and temperature (pH < 2.5, 0°C) before pepsin digestion and measurement of the relative mass increase from deuterium uptake determined by MS/MS. The sequence coverage obtained from the Wuhan spike protein was 64% when in the presence of ICO-hu23, with a peptide-per-residue redundancy of 2.74, and 68% / 2.92 (coverage / redundancy) when in the presence of ICO-hu104. Sequence coverage (and redundancy) for Omicron and Delta variants were similar when complexed with ICO-hu104m at 62% (2.07) and 65% (2.92) respectively **(Supplementary Figs 1A-D)**. Regions of decreased uptake lie within the NTD and RBD, in addition to the S2 helices and helical hinges **(Figure 2A-D)**, corresponding to a reduction in conformational freedom, which agrees with previous Cryo-EM results [14].

**Figure 2.**
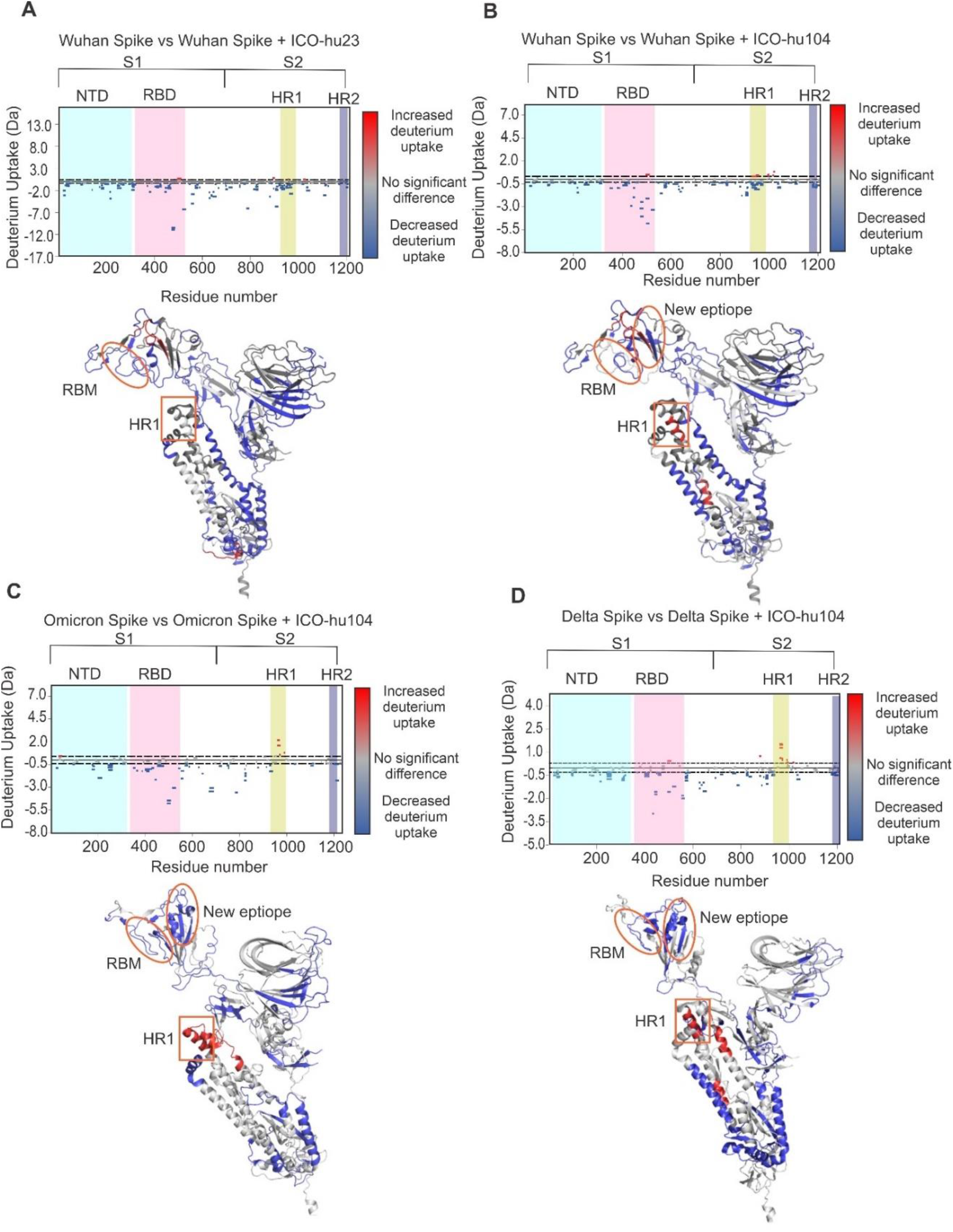
Cumulative Woods plots for (A) Wuhan spike trimer plus ICO-hu23, (B) Wuhan spike trimer plus ICO-104, (C) Omicron spike trimer plus ICO-hu104 and (D) Delta spike trimer plus ICO-104. Peptides with significant increases and decreases in deuterium uptake are coloured on the corresponding spike structures below each of the Woods plots in red and blue respectively. The RBM, new epitope and HR1 are highlighted on the structures by orange boxes. Deuteros was used to produce the Woods plots with an applied confidence interval of 98%.

ICO-hu104 exhibited the same ‘alternate’ epitope for all variants, located between residues 433-443, corresponding to our predictions based on the interaction with hACE2 and agreeing exactly with the epitope identified when using the RBD-only constructs. Binding at this site appeared to invoke a unique allosteric response, whereby an increase in deuterium uptake in the S2 coiled-coil region between, specifically in the HR1 region was observed **(Figures 2A-D)**. Given this unique profile compared to ICO-hu23, and the distinct location of the ICO-hu104 epitope, this could be indicative of the antibody preferentially binding to the ‘up’ RBD position, making residues 433-443 accessible for ICO-hu104 binding. It is also possible that in solution, the spike trimer was not fully occupied by ICO-hu104, diminishing more subtle structural changes and leaving only the most significantly impacted regions to be observed [27]. Nevertheless, HDX-MS reveals a possible neutralization mechanism achieved through allosteric regulation of HR1, which is divergent from its predecessor ICO-hu23.

### Computational modelling reveals allosteric coupling between ICO-hu104 epitope

To explore the allosteric influences of both ICO-hu23 and ICO-hu104 in spike trimer variant structures (Wuhan and Omicron), we applied rigidity transmission allostery (RTA) algorithms. This computational approach furnishes a mechanistic model of allosteric communication by predicting is local perturbations of rigidity propagate from one site and modify rigidity and conformational degrees of freedom at distant sites across the complex. By perturbing the rigidity of ICO-hu23 epitope on Wuhan spike, we found that allosteric communication was more localised within the RBD domain **(Figure 3A)**. Intriguingly, perturbation of the ICO-hu104 epitope on Wuhan spike resulted in allosteric effects to propagate into the S2 coiled-coil region, with stronger allosteric effects in S2 coiled-coil region being observed for the Omicron spike **(Figures 3B and C)**. These computational predictions are in close agreement with experimental HDX observations, validating that ICO-hu104 binding on spike promotes strong allosteric and conformational coupling to propagate the S2 coiled-coil region.

**Figure 3.**
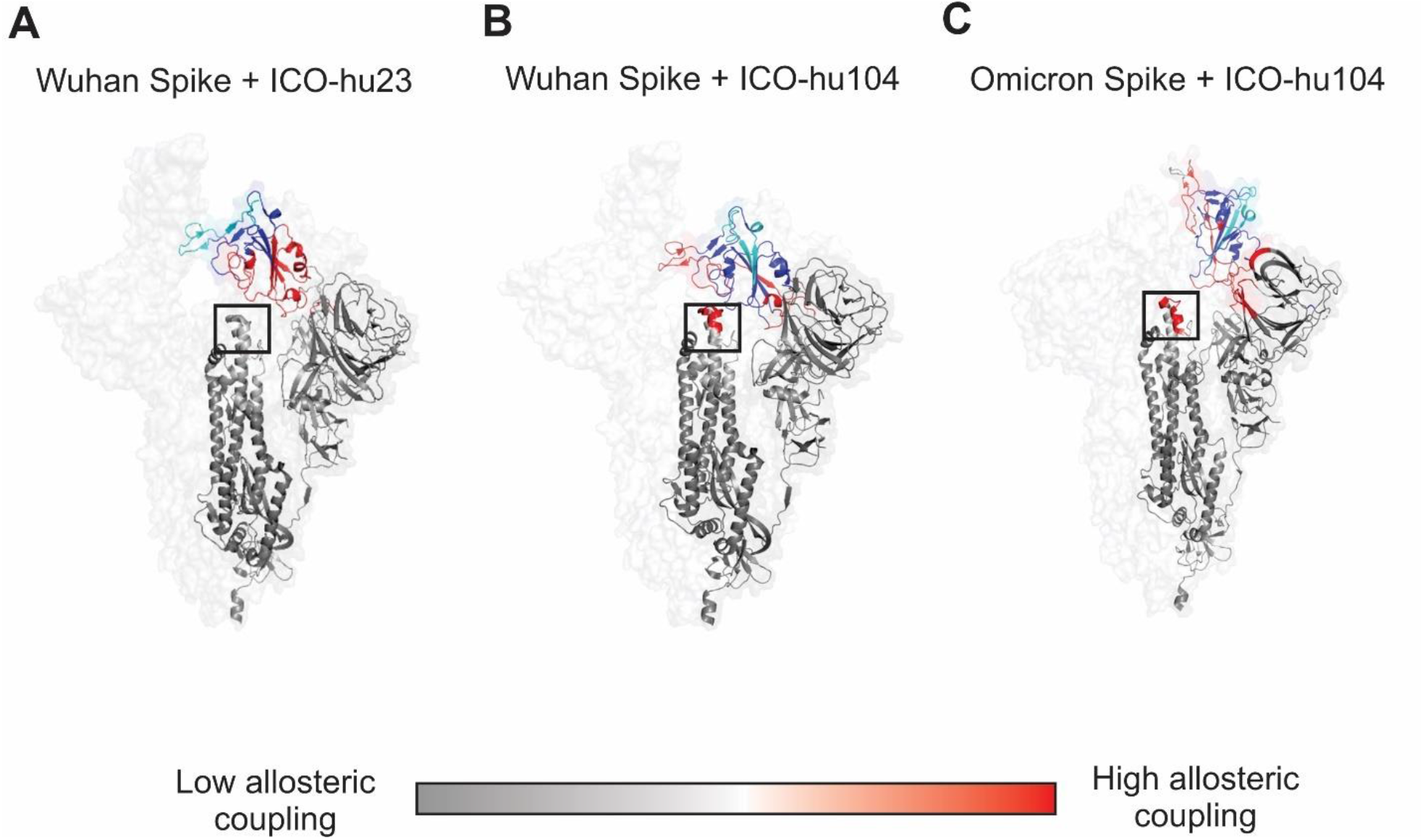
Allosteric effects of ICO-hu23 and ICO-hu104 on Wuhan (PDB 7DDN) and Omicron (PDB 7TGW) spike trimers. (A) RTA allostery predictions are visualised by perturbing the rigidity of the epitope (light blue) of ICO-hu23 on Wuhan spike, (B) ICO-hu104 on Omicron spike and (C) ICO-hu104 on Omicron spike. Dark blue regions are ignored from allostery calculations due to high proximity to the epitopes. Dark red regions indicated high allosteric coupling. S2 coiled-coil region is indicated in the black square.

## Discussion

The rapid development of antibody-based therapeutics is essential for combatting viruses such as SARS-CoV-2, particularly in the early stages of the pandemic, but does present some distinct challenges, including antigenic drift which can quickly render therapies ineffective [28]. Here using HDX-MS, we have successfully characterised allosteric perturbations upon binding of ICO-hu104 in the context of the full-length spike protein against Wuhan, Omicron and Delta variants. Binding of ICO-hu104 to the full-length spike protein distally to the hACE2 binding site in VoC appeared to increase the flexibility within the HR1 region of S2, suggestive of helix destabilisation within the coiled-coil region. Moreover, computational modelling supported that allosteric coupling between the first-generation antibody (ICO-hu23) and the S2 region is minimal, while binding at the ICO-hu104 RBD epitope induces significant allosteric coupling. Together, this suggests a neutralization mechanism which prevents S2-mediated fusion to the host cell.

The currently accepted mechanism of S2-mediated fusion involves exposure of HR1 and HR2 upon insertion of hydrophobic fusion peptides in the S2 homotrimer [29]. Subsequently, a six helical coiled coil is formed as a result of HR2 peptides arranging in an antiparallel fashion and folding into the surface of the HR1 trimeric α-helical core; thus, ensuring close contact for membrane fusion [30]. Increased deuterium uptake within this region indicates weakening of the intra-/inter-helical hydrogen bonding network, increasing flexibility. This ‘unravelling’ of helices that normally extend during S2-mediated fusion suggests that binding at the ICO-hu104 epitope interferes specifically with this process. A number of SARS-CoV-2 neutralizing therapies have been proposed to specifically target the S2 region [31]. Indeed, both HR1 and HR2 within the S2 region have been identified to be highly conserved in VoC [32], therefore making them attractive targets for therapeutics to circumvent mutational changes within the RBD [33]. Specifically, *via* the fusion peptide region or alternatively, small molecule therapies have been proposed to disrupt the region between HR1 and HR2 however, little progress has been made towards characterising their binding sites [34]. Nevertheless, identification of novel neutralization mechanisms governed by allosteric control broadens our understanding of alternative neutralization pathways which could help combat immune evasion.

Unravelling allosteric perturbations within the S2 region of SARS-CoV-2 upon binding of ICO-hu104 is essential to elucidate its neutralization mechanism. Together, these insights can inform on strategies to quickly characterise candidate therapeutic antibodies, and to support the selection of those with favourable neutralization mechanism to overcome antigenic drift. Furthermore, the application of HDX-MS and ability to uncover allosteric perturbations quickly can open new avenues for antibody design, uncovering novel mechanisms of action which can complement existing therapies.

## Accession numbers

Severe acute respiratory syndrome coronavirus 2 (2019-nCoV) (SARS-CoV-2) Spike glycoprotein is associated with the UniProt accession code P0DTC2. The Wuhan SARS-CoV-2 Spike glycoprotein is associated with the PDB accession code 7DDN, the Omicron SARS-CoV-2 Spike glycoprotein is associated with the PDB accession code 7TGW, and the Delta SARS-CoV-2 Spike glycoprotein is associated with the PDB accession code 7ZJL.

## CRedIT authorship contribution statement

**Alice Colyer:** Writing-original draft, writing-review and editing, validation, visualisation.

**Esther Wolf:** Writing-review and editing, conceptualization, data curation, formal analysis, investigation, methodology, validation, visualisation.

**Cristina Lento:** Writing-review and editing, methodology product administration, resources, software, supervision

**Mart Ustav Jr**.: Writing-review and editing, conceptualization, funding acquisition, methodology, product administration, resources.

**Adnan Sljoka:** Writing-review and editing, conceptualization, data curation, formal analysis, investigation, methodology, validation, visualisation.

**Derek J. Wilson:** Writing-review and editing, conceptualization, funding acquisition, methodology, product administration, resources, software, supervision.

## Acknowledgements

We would like to thank Dr. Andrew James and Dr. Dominic Narang for helpful discussions.

