## Supplementary Figure 1A-D for "Allostery links hACE2 binding, pan-variant neutralisation and helical extension in the SARS-CoV-2 Spike protein"

***Contributed equally**

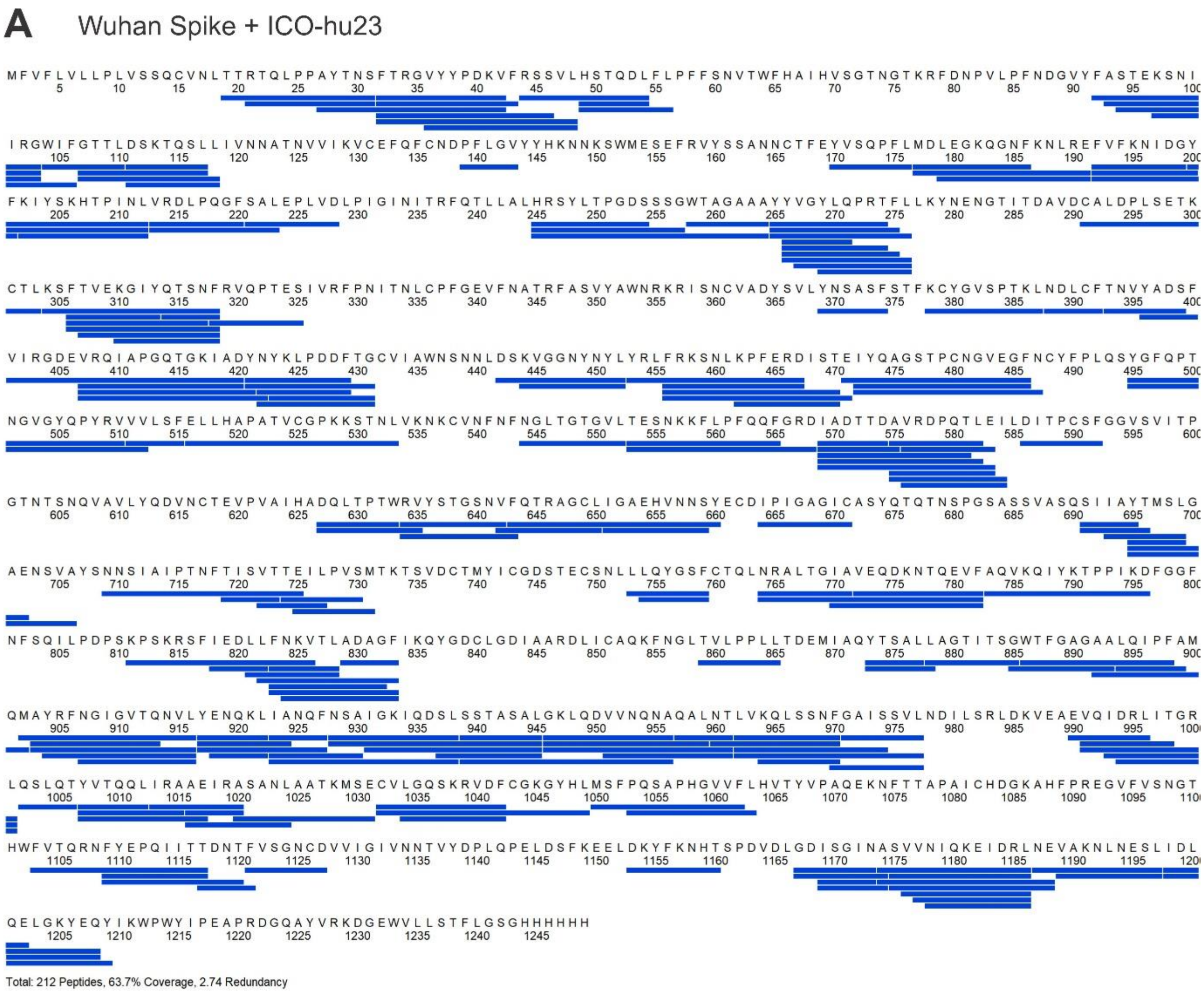


**Supplementary Figure 1** (A) Sequence coverage map of full-length Wuhan Spike coupled to ICO-hu23. Sequence coverage after peptide assignment was 64% with a redundancy of 2.74.

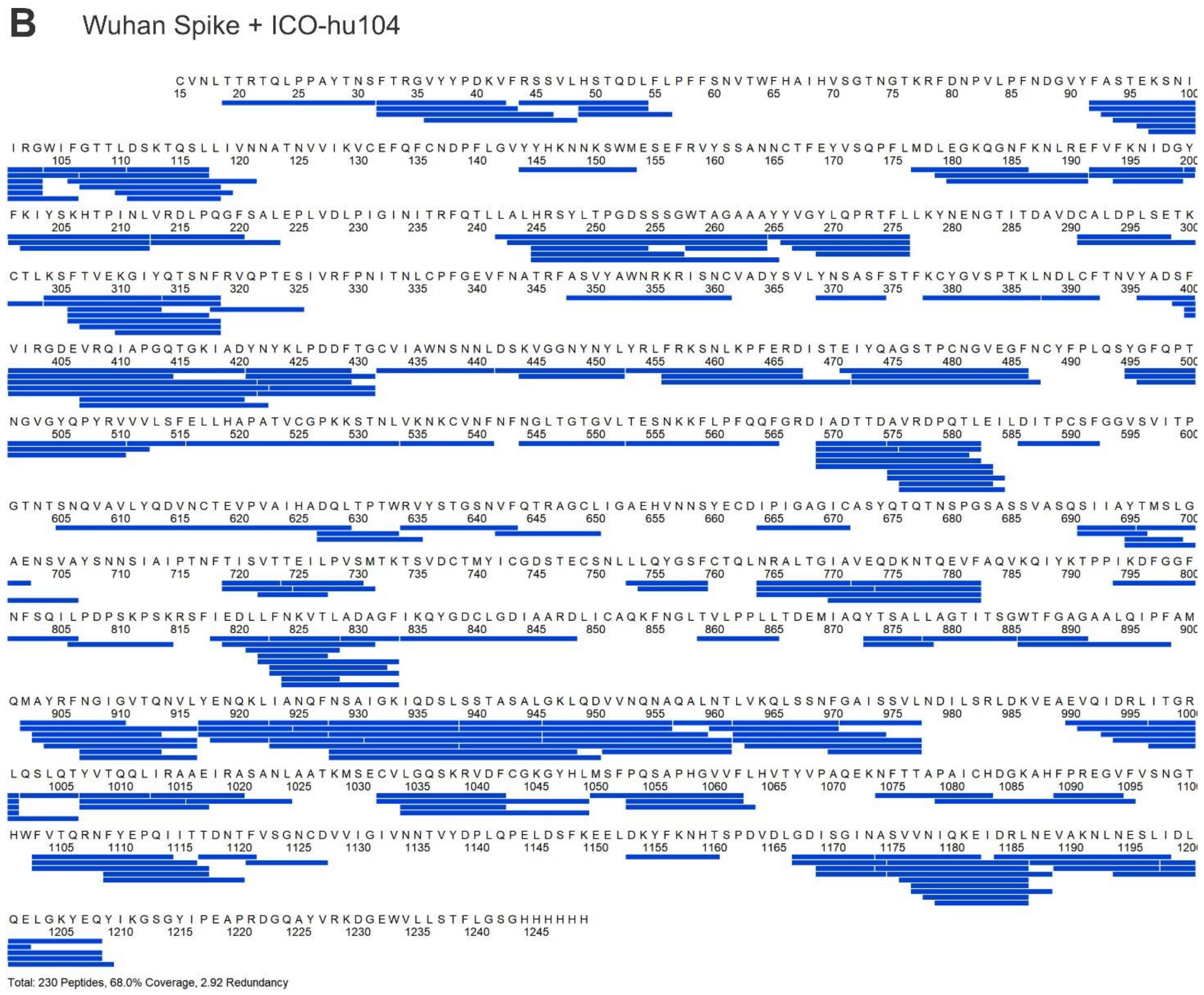


**Supplementary Figure 1** (B) Sequence coverage map of full-length Wuhan Spike coupled to ICO-hu104. Sequence coverage after peptide assignment was 68% with a redundancy of 2.92.

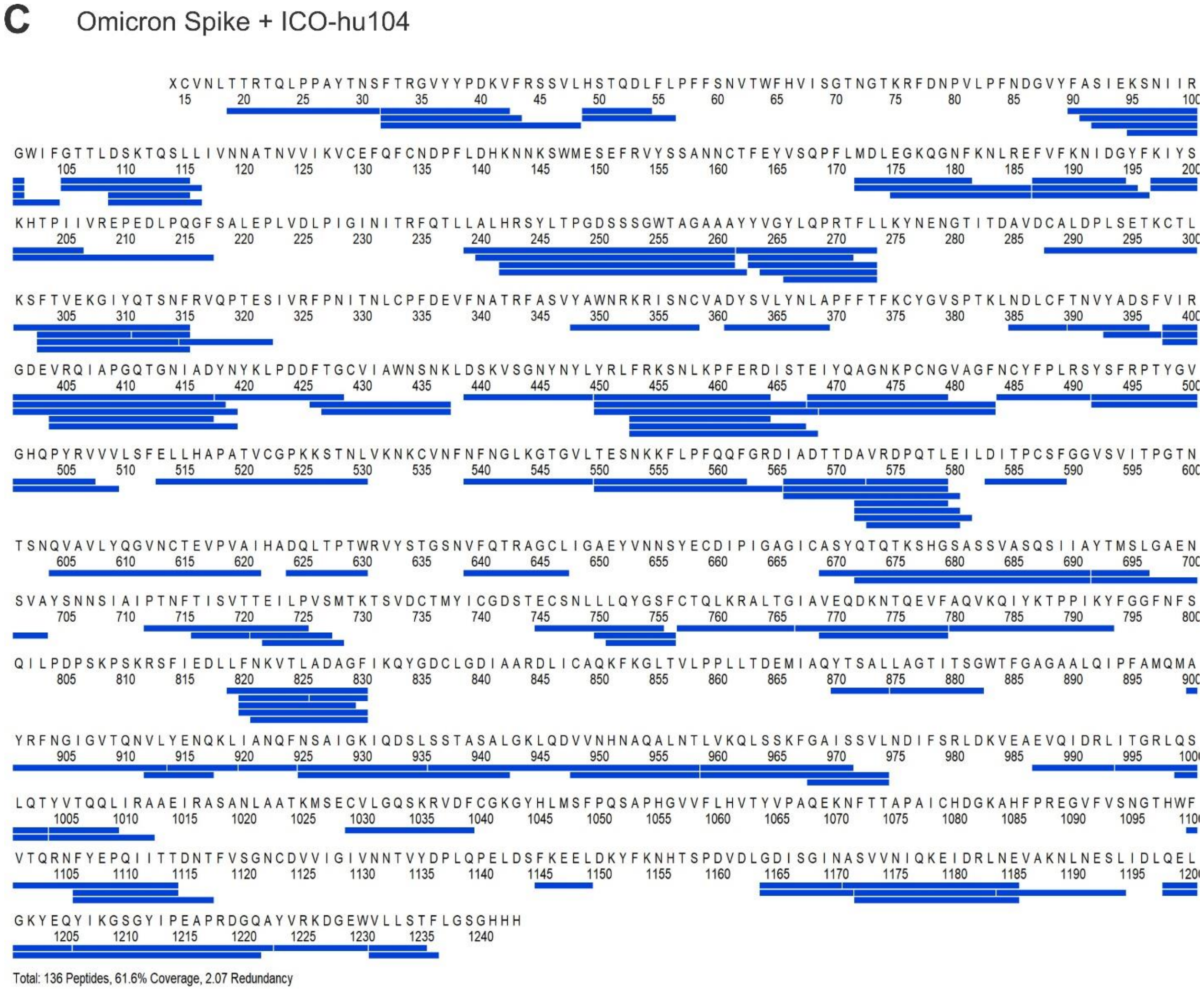


**Supplementary Figure 1** (C) Sequence coverage map of full-length Omicron Spike coupled to ICO-hu104. Sequence coverage after peptide assignment was 62% with a redundancy of 2.07.

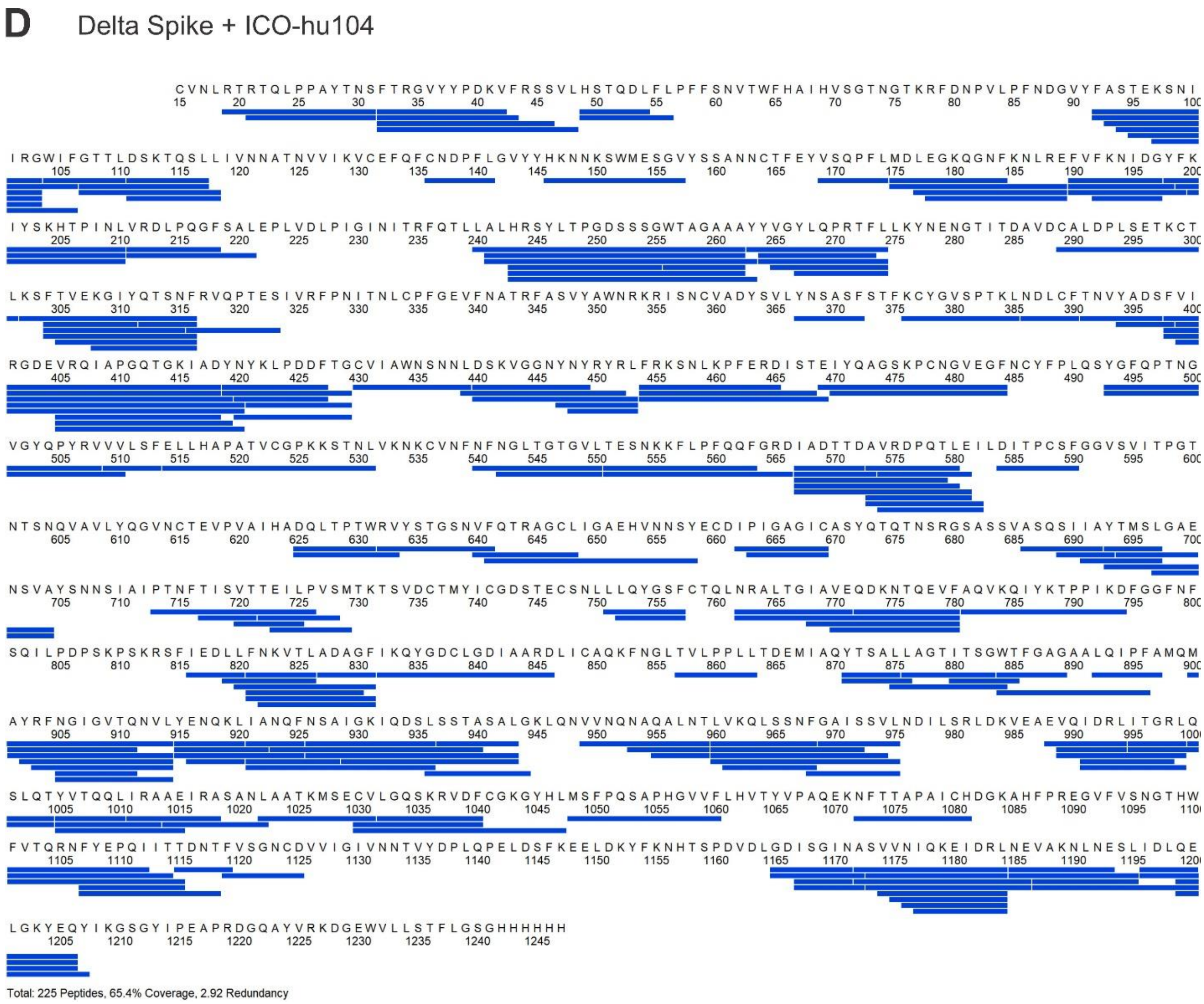


**Supplementary Figure 1** (D) Sequence coverage map of full-length Delta Spike coupled to ICO-hu104. Sequence coverage after peptide assignment was 65% with a redundancy of 2.92.
